# Why are education, socioeconomic position and intelligence genetically correlated?

**DOI:** 10.1101/630426

**Authors:** Tim T Morris, Neil M Davies, Gibran Hemani, George Davey Smith

## Abstract

Genetic associations and correlations are perceived as confirmation that genotype influences one or more phenotypes respectively. However, genetic correlations can arise from non-biological or indirect mechanisms including population stratification, dynastic effects, and assortative mating. In this paper, we outline these mechanisms and demonstrate available tools and analytic methods that can be used to assess their presence in estimates of genetic correlations and genetic associations. Using educational attainment and parental socioeconomic position data as an exemplar, we demonstrate that both heritability and genetic correlation estimates may be inflated by these indirect mechanisms. The results highlight the limitations of between-individual estimates obtained from samples of unrelated individuals and the potential value of family-based studies. Use of the highlighted tools in combination with within-sibling or mother-father-offspring trio data may offer researchers greater opportunity to explore the underlying mechanisms behind genetic associations and correlations and identify the underlying causes of estimate inflation.

## Introduction

Genetic associations with phenotypes can be used to interrogate causal relationships, provide insight into the genetic architecture of complex traits, how relationships between phenotypes operate, and indicate future intervention strategies. A persons own educational attainment and their parents occupational position are two of the strongest determinants of health and social outcomes throughout the lifecourse.^1–5^ Education and occupational position are conceptually viewed as subcategories of broader socioeconomic position (SEP),^6,7^ are strongly correlated^8–10^ and highly heritable.^2,5,11–13^ Furthermore, education has been demonstrated to have a complex genetic architecture characterised by high polygenicity,^14,15^ and this is likely to be the case for other complex social traits. The extent to which genetic factors influence educational attainment and SEP independently and jointly has important implications for social policy to reduce social and educational inequalities throughout the lifecourse. Misinterpretation of genetic associations could lead to misleading conclusions and even potentially harmful interventions. Genetic associations are sensitive to a range of assumptions, and bias can be introduced by violations to these assumptions. In this paper we outline alternative explanations for estimated genetic associations, demonstrate some of the available tools to assess these biases, indicate future study designs that can bring further clarity to this area of research, and provide a framework for addressing questions that determine the source of genetic associations. Studies with a genetic focus that study complex social phenomena^16,17^ may be particularly susceptible to these biases and we therefore focus on the growing area of sociogenomics, but our findings are relevant to the broader genetic literature.

Educational attainment is one of the most heavily studied social phenotypes in genetic studies.^14^ The heritability of education has been estimated at 40%^2^ for years of education and ~60% for test score attainment.^18–21^ There is evidence of genetic correlation between education and other indicators of SEP. Genetic correlation refers to the correlation of the genetic contributions to two phenotypes and is commonly used to demonstrate the presence of shared genetic architecture. Genetic associations are estimated using individual or GWAS summary data with methods such as GREML^22^ and LD Score regression^23,24^ respectively. A review of the relative merits of these methods is provided by Ni et al.^25^ Social class and educational attainment have been estimated to be highly genetically correlated at 0.87 (0.32),^5^ 1.02^i^ (0.25),^11^ and 0.48 (0.02)^12^ using GREML. The implicit understanding of a genetic correlation is that there are shared biological mechanisms within an individual that influence both their education and their SEP, that is, correlation between causal genetic variants. One strong phenotypic candidate that may underlie this relationship is cognitive ability,^11,26,27^ itself highly heritable^12,13,18^ and with strong correlations with educational attainment and other measures of SEP.^1,8,28–32^ Genetic correlations between cognitive ability and educational attainment have been estimated at 0.65 (0.02)^12^ and 0.63 (0.03).^18^ Genetic correlations between cognitive ability and measures of socioeconomic position have been estimated at 0.40 (0.02)^12^ using neighbourhood social deprivation and 1.00 (0.47)^13^ using a composite measure of parents’ occupation and education. While education, occupation and deprivation are components of broader socioeconomic position, these non-zero genetic correlations between phenotypes have been interpreted as evidence that complex social outcomes are influenced by overlapping genetic factors.

However, estimates from methods based on unrelated individuals are sensitive to bias introduced through violations of assumptions^33,34^. That is, genetic associations can arise from confounding factors as well as biological effects. Apparent genotype-phenotype associations can arise from non-biological, environmental or indirect mechanisms instead of a causal biological mechanism in which genetic variation influences both phenotypes, changing the interpretation of large genetic associations. Possible alternative mechanisms which can induce genetic associations between phenotypes are demonstrated in Figure 1 and include horizontal pleiotropy,^35^ population stratification,^36^ dynastic effects,^37^ and assortative mating.^38,39^

**Figure 1:**
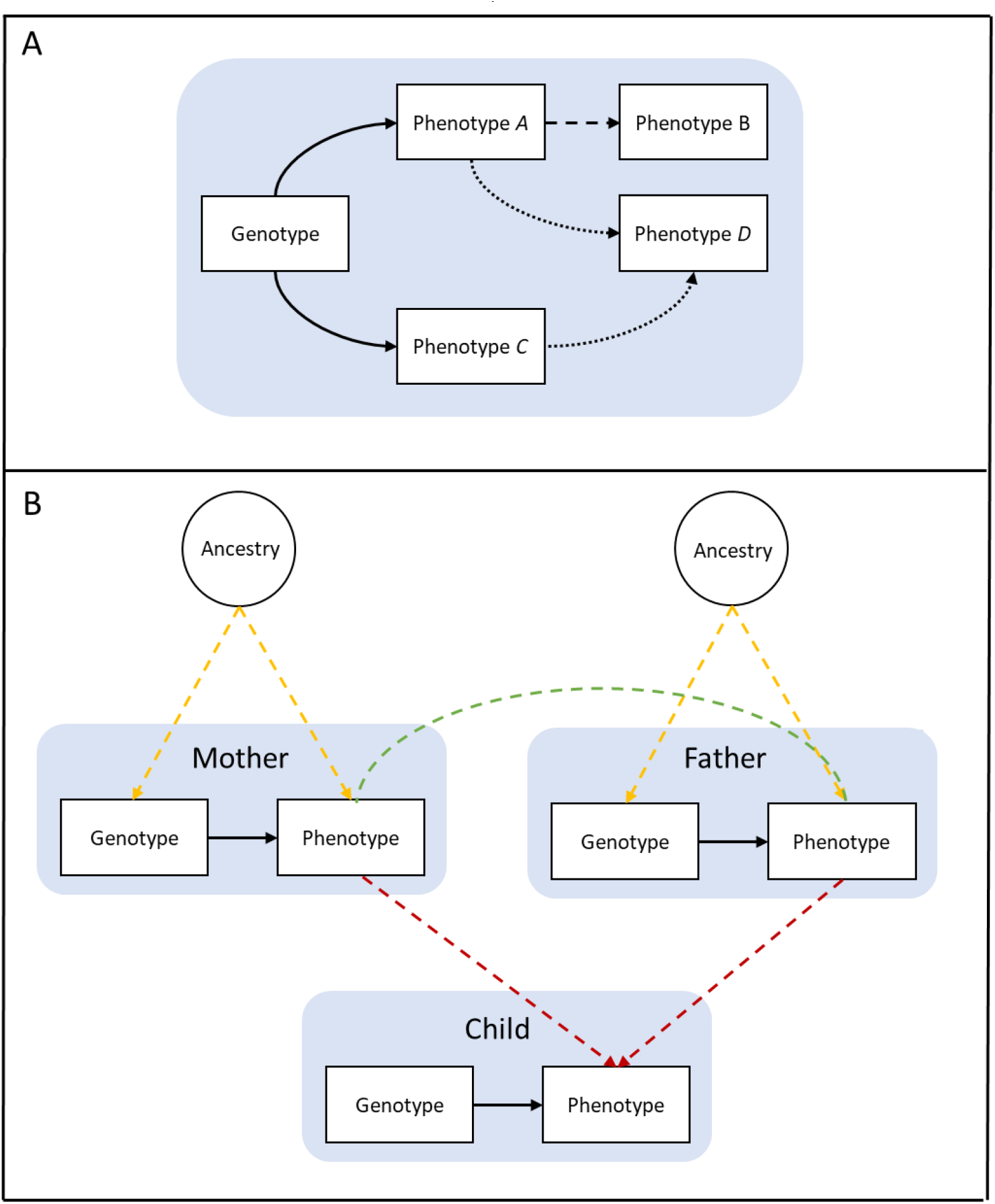
Causal models of structures underlying genetic associations. Panel A: ‘Direct’ causal effect where genetic expression operates directly on phenotype *A* (solid line); ‘Indirect’ causal effect where genetic expression operates on phenotype *B* only through phenotype *A* (dashed line), referred to as vertical pleiotropy; ‘Direct’ horizontal pleiotropy where genetic expression simultaneously influences phenotypes *A* and *C* (solid lines); ‘Indirect’ horizontal pleiotropy where genetic expression operates on phenotype *D* through phenotypes *A* and *C* simultaneously (dotted lines). Panel B: Population stratification due to ancestral differences (yellow lines); Dynastic effects (red lines); Assortative mating (green line). Arrows represent direction of effect; nondirected lines represent simultaneous assortment.

## The causes of genetic associations: mechanisms and tools to assess these

There are various structures by which univariate and bivariate genetic associations can arise (Box 1). These include causal genetic effects; population structure; dynastic effects; and assortative mating. Causal genetic effects are often the interest of researchers. They can operate ‘directly’ whereby genotypic expression exerts an influence on a phenotype (Figure 1a, solid lines). In the absence of bias and confounding, univariate genetic association between the SNP and trait can provide evidence of causal effect of genotype on phenotype(s). ‘Indirect’ causal genetic effects may also operate whereby genotype influences a phenotype *only through* another phenotype (Figure 1a, dashed line), referred to as vertical pleiotropy. Vertical pleiotropy is the mechanism of causation targeted in Mendelian Randomization studies to estimate causal effects of phenotype *A* on outcome *B*, with genotype *G* instrumenting for *A*. Provided that the instrumental variable assumptions hold, Mendelian Randomization can be used to estimate the causal effect of phenotypes on outcomes. However, genotype may influence the outcome through multiple pathways simultaneously (Figure 1a, dotted lines), which is referred to as horizontal pleiotropy. Horizontal pleiotropy is a concern for Mendelian Randomization studies (for a detailed discussion see ^35^), whereas genetic correlations typically reflect the correlation in effects of common SNPs across the entire genome on two or more phenotypes. These correlations can be due to either the causal effects of genetic variants or pleiotropic effects, or any other source of bias. Genetic correlations and Mendelian randomization use different estimators; genetic correlations use genetic variation across the entire genome, while Mendelian randomization typically uses only very few genetic variants for which there is strong evidence they associate with the exposure. Genetic correlation between two phenotypes is normally necessary but not sufficient to indicate a causal relationship (for a discussion of genetic correlation and MR see ^40,41^). However, genetic correlations do not provide evidence for directionality in associations.

One such biasing mechanism which may explain genetic associations is population stratification (Figure 1b, yellow lines), which refers to systematic differences in allele frequencies between subpopulations arising from ancestral differences. Population stratification can induce spurious associations between genotype and phenotype even where a causal relationship is absent. Because of the historical importance of migration in driving genetic differences, population stratification often appears as geographical structure.^36,42^ It is commonly controlled for by adjusting for principal components of genotype within the sample. Principal components are designed to capture common differences between sub-populations in allele frequencies, thus removing bias caused by population stratification. A recent study demonstrated that geographical structure remains even after controlling for the first 100 principal components in UK Biobank, far in excess of the 10 or 20 components commonly controlled for.^43^ If residual population structure associates with two phenotypes, then genetic correlation will spuriously demonstrate apparent shared genetic effects across the genome. Given the non-random complexity of migration and mating over many generations it is unlikely that population stratification will be fully controlled for in samples of unrelated individuals. While it is not possible to prove that adjusting for principal components has controlled for all differences within the sample, one way to assess the impact of population stratification is to compare estimates obtained from unadjusted models and models that adjust for principal components. Attenuation in estimated effect sizes after principal component adjustment provides support for population stratification bias, and the extent of this bias may be gauged by the extent of attenuation. However, in regional studies this may not be sufficient.^43^ Between-sibling and within family study designs may also be used as they are robust to population stratification: because siblings share the same genotype source, differences between them cannot be attributed to population stratification. Such studies require very large samples due to lower phenotypic and genotypic variation between siblings than unrelated individuals.

Genetic associations can also be induced/biased by dynastic effects (Figure 1, path red lines) where inherited genetic variants operate indirectly on offspring phenotype via their expression in the parents’ phenotype. For example, if education associated variants at the parental generation contribute to the creation of education enriching environments through the provision of books in the household, then offspring who inherit education associated variants are also more likely to inherit education associated environments. This is a form gene-environment correlation. Thus, social or ‘environmental’ transmission effects can affect SNP-phenotype associations, which can make them biased estimates of the causal effect of the SNP on the phenotype. Where parental genotype influences the same phenotype in both parents and offspring this can be thought of as a double contribution or ‘double inheritance’ of genotype. It is possible that this explains the relatively low estimates of the contribution of the shared environment from twin studies. There is a large body of evidence suggesting that social phenotypes such as educational attainment and socioeconomic position are socially transmitted across generations,^44–46^ and it is possible that genetic associations with these phenotypes will be affected by dynastic effects. The presence of dynastic effects can be tested with data on mother-father-offspring trios or siblings.^47^ Using polygenic scores, the raw association between offspring genotype and phenotype can be compared with its association when adjusted for mother and father genotype. Attenuation of the raw association and direct (conditional) association between parental genotype and offspring phenotype supports an indirect effect of parental genotype on offspring phenotype and therefore the presence of dynastic effects. It is also possible to use non-transmitted parental variants to create a ‘genetic nurture’ polygenic score.^37^ Because non-transmitted variants can only influence offspring phenotype indirectly, association between a non-transmitted score and offspring phenotype supports dynastic effects. Relatedness disequilibrium regression, which exploits variation in relatedness due to random segregation, can also be used to estimate the bias induced to heritability estimates by environmental effects.^34^ The above methods all require genetic data on mother-father-offspring trios.

Assortative mating (Figure 1, path green line), or social homogamy, may also induce genetic associations between phenotypes. Assortative mating refers to the non-random pairing of spouses across the population due to mate selection based on phenotypic characteristics. There is evidence for assortative mating on a range of phenotypes including educational attainment and socioeconomic position.^38,39,48–50^ Where phenotypes that are selected on have a genetic component, then assortative mating will lead to spouses being more genetically similar to each other than to randomly selected individuals from the population. That is, phenotypic assortative mating across the population leads to an increase in the likelihood of people mating with partners who are more genetically similar. While random mating would ensure even distribution of allele frequencies at the population level, assortative mating leads to systematic differences in allele frequencies (population stratification) and subsequent deviations from Hardy-Weinberg Equilibrium over generations.^38^ Genotype-phenotype associations in subsequent generations can therefore be biased by phenotypic assortment. Assortative mating will lead to a disproportionate enrichment or depletion of education alleles within spouse couples and increased homozygosity in offspring across a population. That is, offspring of parents with high educational attainment will be more likely to have education increasing alleles than offspring of parents with low educational attainment. This will inflate heritability estimates due to the increased genetic variance in the population.^51^ If spouses sort on different traits (i.e. cross trait assortative mating) then assortment can also induce genetic correlations between traits in offspring.^52^ Assortative mating can lead to population stratification if it is sub-population specific,^53^ and to disproportionate inheritance of the environment in addition to genotype if dynastic effects exist. Assortative mating can therefore bias estimates of genetic correlation and Mendelian randomization.^54^

Statistical methods to estimate genetic associations from unrelated individuals assume no population stratification, dynastic effects or assortative mating. Where any of these structures exist and are insufficiently controlled for, estimates of genetic associations will be inflated due to these hidden correlations in the data and the confounding effects will be incorrectly attributed to genetic effects.^48^ Using the example of educational attainment and socioeconomic position in a UK birth cohort, we first present univariate and bivariate genetic associations and then investigate the presence of the above biasing mechanisms in our results.

### Box 1: Structures which can induce genetic associations.

Causal genetic effects; population stratification; dynastic effects and assortative mating.

#### 1. Causal genetic effects

Causal genetic associations operate where genetic variation has a causal effect on the phenotype(s). GWAS studies usually aim to estimate causal genetic effects. Causal effects may be investigated for a single phenotype (univariate genetic association) or pairs of phenotypes (bivariate genetic association or genetic correlation).

#### 2. Population stratification

Population stratification refers to the systematic difference of allele frequencies across subpopulations. This arises from ancestry differences due to non-random mating and subsequent genetic drift of allele frequencies between sub-population groups, historically caused by geographic and physical boundaries. If phenotypes also differ systematically between subpopulations, population stratification can lead to genotype-phenotype associations despite no causal relationship between the genotype and the phenotype.

#### 3. Dynastic effects

Further to influencing offspring phenotype through genetic inheritance, parental genotype can indirectly influence offspring phenotype through its expression on parental phenotype. Where this occurs, offspring may inherit both phenotype associated variants and phenotype associated environments from parents, leading to upwards bias in genetic associations.^37^ For example, genetic variants positively associated with education in the parent’s generation may lead to the creation of educational rich environments (such as an increase in books in the household) which will have a positive impact upon the child’s educational attainment. Dynastic effects refer to this ‘inheritance’ of environment in addition to genotype.

#### 4. Assortative mating

Assortative mating refers to the process by which spouses select each other based upon certain phenotypes. If these phenotypes have a genotypic component, then phenotypic selection induces greater genetic similarity between spouses than in the general population. This can induce correlation between phenotype and genotype in subsequent generations and lead to overestimation of the contribution of genotype to phenotypic associations because genetic associations arise due to confounding by phenotypic assortment.^51,52,54^ While offspring inheritance of genotype is random conditional upon parent’s genotype, assortative mating induces non-random inheritance patterns across groups based on phenotype.

## Results

Table 1 displays the raw variables used in the analyses. Compared to participants who were excluded due to missing data, those included in the analyses had higher attainment at each stage of education, higher cognitive ability as measured at age 8, and came from higher SEP families as measured on both linear and binary.

**Table 1:** Descriptive statistics for raw variables.

|  | Included |  |  | Excluded |  |  | *p* value |
| --- | --- | --- | --- | --- | --- | --- | --- |
|  | n | mean | SD | n | mean | SD |  |
| Age 11 score | 6,132 | 28.04 | 3.85 | 4,976 | 26.74 | 4.20 | <0.001 |
| Age 14 score | 4,960 | 35.97 | 6.19 | 4,055 | 33.65 | 6.56 | <0.001 |
| Age 16 score | 6,518 | 39.89 | 9.48 | 5,228 | 36.46 | 10.09 | <0.001 |
| Cognitive ability | 5,295 | 105.07 | 16.36 | 1,953 | 101.34 | 16.67 | <0.001 |
| Linear SEP | 6,418 | 59.35 | 13.42 | 4,524 | 55.49 | 13.49 | <0.001 |
| Binary SEP | n | % |  | n | % |  | <0.001 |
| High | 2,713 | 59.53 |  | 2,302 | 48.77 |  |  |
| Low | 3,990 | 40.47 |  | 2,418 | 51.23 |  |  |

SEP, socioeconomic position.

### Univariate phenotypic heritability

SNP heritability of educational attainment increases with age from 44.7% (95%CI: 32.7 to 56.6) at age 11 to 52.5% (95%CI: 37.8 to 67.0) at age 14 and 61.2% (95%CI: 50.2 to 72.2) at age 16 (Figure 2a). The heritability of cognitive ability is estimated at 45.2% (95% CI: 33.0 to 57.6), while the heritability of socioeconomic position is estimated to be higher for the linear measure (53.0%; 95% CI: 42.9 to 63.0) than the binary measure (33.9%; 95% CI: 24.2 to 43.5). Interestingly, the heritability for the linear measure of socioeconomic position is comparable to that of educational attainment.

**Figure 2:**
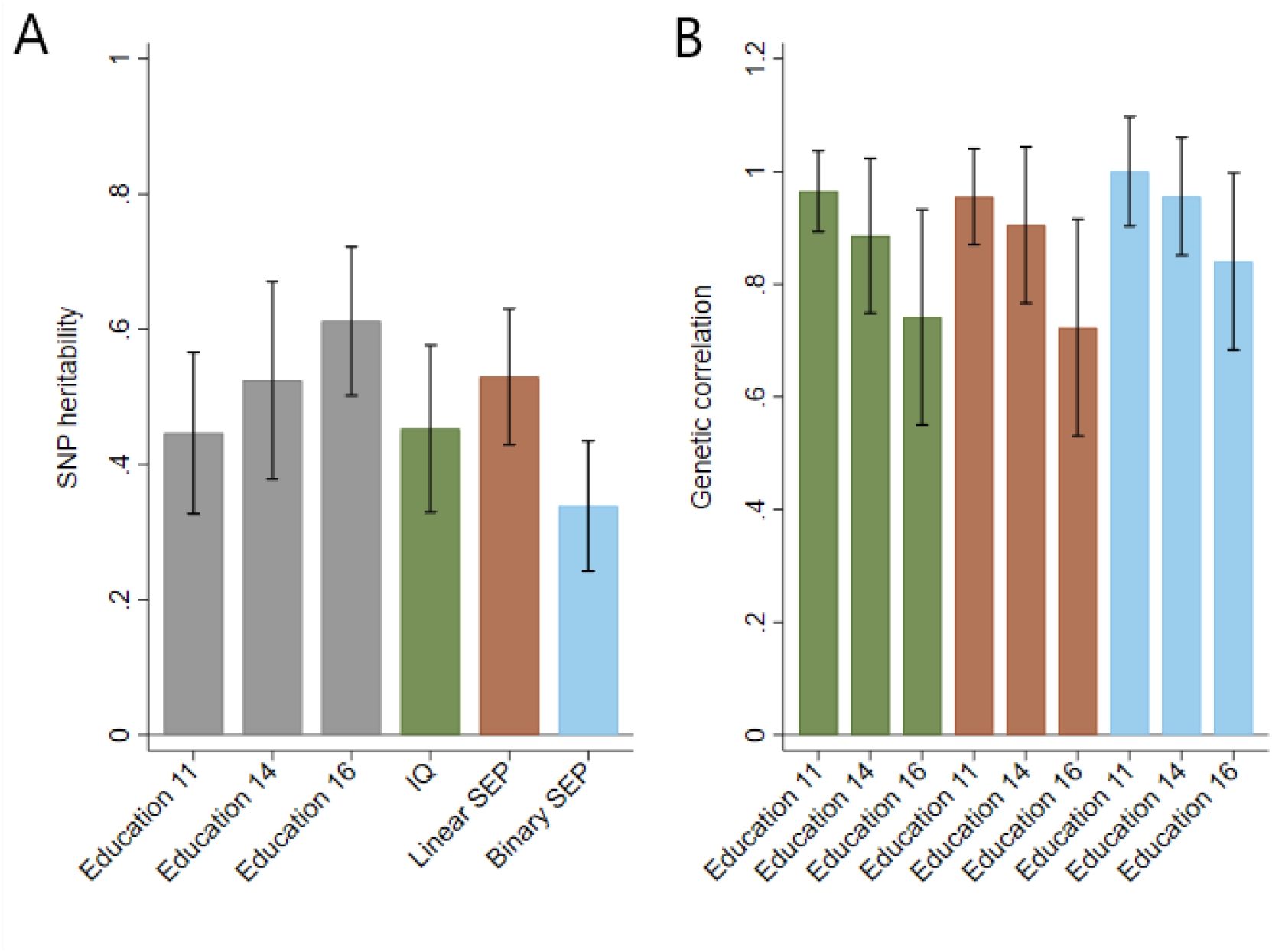
SNP heritability and genetic correlations between phenotypes. Panel A: Grey bars represent educational attainment measured at ages 11, 14 and 16; green bar represents cognitive ability measured at age 8; orange bar represents linear socioeconomic position; blue bar represents binary socioeconomic position. Panel B: Green bars represent genetic correlations between educational attainment at ages 11, 14 and 16 with cognitive ability measured at age 8; orange bars represent genetic correlations between educational attainment at ages 11, 14 and 16 with linear socioeconomic position; blue bars represent genetic correlations between educational attainment at ages 11, 14 and 16 with binary socioeconomic position. See Tables 1 and 2 in supplementary material for full results.

### Genetic correlation

Genetic correlations between educational attainment and cognitive ability are high and persist throughout childhood within the range of 0.96 to 1 (Figure 2b). This suggests that many SNPs which associate with educational attainment also associate with intelligence. Genetic correlations between educational attainment and socioeconomic position are also high: For the linear measure they range from 0.89 (95% CI: 0.75 to 1.02^ii^) to 0.96 (95% CI: 0.85 to 1.06), and for the binary measure from 0.76 (95% CI: 0.57 to 0.95) to 0.87 (95% CI: 0.71 to 1.04). The genetic correlations suggest that many SNPs which associate with educational attainment also associate with family socioeconomic position. To check that our results were not driven by genotyping or imputation method we repeated our analyses using data imputed to 1000 genomes. The results were consistent using each method (Tables 1–4 in supplementary material).

The polygenic score for educational attainment explains between 3.6 and 5.1% of the variation in educational attainment, 3.0% in cognitive ability and the linear measure of socioeconomic position, and 1.6% in the binary measure of socioeconomic position (Figure 3). That the polygenic score explains a similar amount of variation in the linear measure of socioeconomic position as educational attainment suggests a modest amount of pleiotropy in the variants used in the score. This underscores the high genetic correlations observed above.

**Figure 3:**
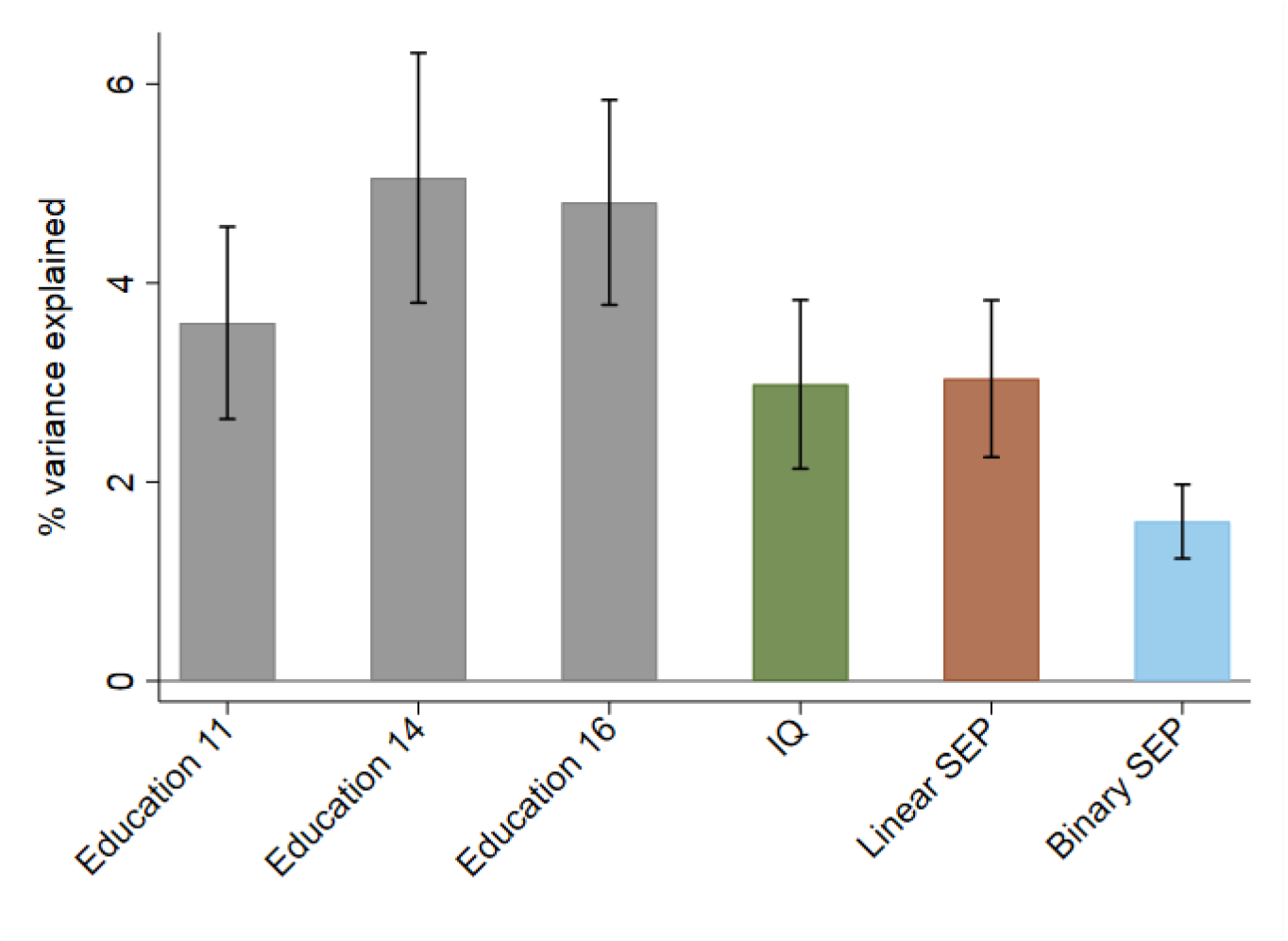
Variance explained by the EA score. Variance explained in each phenotype by the educational attainment polygenic score constructed from genomewide significant variants. Standard errors were obtained through bootstrapping with 1000 replications. Grey bars represent educational attainment measured at ages 11, 14 and 16; green bar represents cognitive ability measured at age 8; orange bar represents linear socioeconomic position; blue bar represents binary socioeconomic position.

### Bivariate SNP heritability

While genetic correlation estimates the correlation between the effects of SNPs on two traits, bivariate heritability estimates the proportion of phenotypic correlation between two traits that can be attributed to coheritability (equation 4 in Materials and methods). The bivariate heritabilities of educational attainment and cognitive ability range from 0.69^iii^ at age 11 to 0.70 at age 14 to 0.85 at age 16. At face value this suggests that over two thirds of the phenotypic similarity between educational attainment and cognitive ability can be explained by shared common genetic variation in our sample. The higher bivariate heritability at age 16 arises from lower phenotypic correlation between educational attainment and cognitive ability at this measurement occasion (Table 5 in supplementary material). The bivariate heritabilities for educational attainment and socioeconomic position are estimated at 1.30 (age 11), 1.27 (age 14), and 1.25 (age 16) for the linear measure, and 1.21 (age 11), 1.14 (age 14), 1.24 (age 16) for the binary measure. These bivariate heritability estimates are greater than one because the coheritability (the numerator in equation 4) is greater than the phenotypic correlation (the denominator in equation 4). While unexpected, these high values are possible because bivariate heritability is estimated using covariance which is not constrained by the values of −1 and 1. Bivariate heritabilities greater than one suggest that univariate heritabilities or genetic correlation have been inflated and therefore that coheritability is overestimated. As such they can provide an indication that the biasing mechanisms outlined above are present. We now investigate how these mechanisms may have biased our estimates.

### Exploring potential biasing mechanisms

#### Population stratification

Comparison of the heritability estimates between unadjusted models and models that adjust for the first 20 principal components of the GRM indicates that bias due to population stratification in our results is likely to be low (Table 2). Estimates of heritability are slightly lower in all PC adjusted models, but these differences are very small and could be due to estimation error. While, it is unlikely that adjustment for the first twenty principal components will have removed all population stratification bias^43^, but the small size of attenuation of our heritability estimates suggests that any remaining population stratification bias is likely to be low.

**Table 2:** Testing for population stratification bias.

|  | Unadjusted | Population stratification adjusted |
| --- | --- | --- |
| EA age 11 | 0.498 (0.056) | 0.451 (0.064) |
| EA age 14 | 0.545 (0.069) | 0.509 (0.079) |
| EA age 16 | 0.607 (0.052) | 0.605 (0.059) |
| Cognitive ability | 0.453 (0.058) | 0.417 (0.066) |
| Linear SEP | 0.568 (0.047) | 0.547 (0.054) |
| Binary SEP | 0.372 (0.046) | 0.337 (0.052) |

EA, educational attainment; SEP, socioeconomic position.

#### Dynastic effects

Table 3 shows associations between the study child’s education polygenic score and their educational attainment at age 16 before and after adjustment for both parents polygenic scores, based on a sample of 1,095 mother-father-offspring trios. In the unadjusted model, a one standard deviation higher educational attainment polygenic score built from all SNPs is associated with a 0.340 (0.028) standard deviation higher attainment at age 16. After adjustment for both parents polygenic scores this is attenuated to 0.223 (0.041), an attenuation of 34.4%. Using a polygenic score built only from the education associated SNPs that reached genomewide significance, the association of the child’s polygenic score and their educational attainment attenuated by 60.5% after adjustment for parental polygenic scores. Furthermore, parent’s education genome-wide polygenic scores remained associated with their child’s education attainment conditional on the child’s polygenic score. There was little evidence that the parents’ polygenic score using genome-wide significant hits was conditionally associated with the offspring’s educational attainment. These results potentially demonstrate the presence of educational dynastic effects or assortative mating. Our negative control analyses of C-reactive protein based on 942 mother-father-offspring trios showed that a one standard deviation higher CRP polygenic score was associated with a 0.219 (0.030) standard deviation higher level of CRP. After adjustment for both parents CRP polygenic scores this is attenuated to 0.192 (0.043), an attenuation of 12.4%. Neither the maternal nor paternal CRP polygenic were associated with offspring phenotypic CRP conditional on offspring CRP polygenic score. These findings are consistent with no dynastic or assortative mating effect for CRP.

**Table 3:** Associations between child and parent polygenic scores with phenotypes (n=1,095 trios).

|  | Unadjusted | Co-adjusted | *p* value for difference | Attenuation (%) |
| --- | --- | --- | --- | --- |
| *Education* |  |  |  |  |
| *Education PGS all SNPs* |  |  |  |  |
| Child PGS | 0.340 (0.028) | 0.223 (0.041) | <0.001 | 34.4 |
| Mother PGS | 0.261 (0.029) | 0.110 (0.036) |  |  |
| Father PGS | 0.232 (0.030) | 0.099 (0.034) |  |  |
| *Education PGS GWAS SNPs* |  |  |  |  |
| Child PGS | 0.129 (0.029) | 0.051 (0.042) | 0.009 | 60.5 |
| Mother PGS | 0.072 (0.030) | 0.035 (0.036) |  |  |
| Father PGS | 0.142 (0.029) | 0.111 (0.036) |  |  |
| *CRP* |  |  |  |  |
| *CRP GWAS SNPs* |  |  |  |  |
| Child PGS | 0.219 (0.033) | 0.192 (0.043) | 0.330 | 12.4 |
| Mother PGS | 0.079 (0.033) | -0.002 (0.037) |  |  |
| Father PGS | 0.144 (0.032) | 0.056 (0.037) |  |  |

PGS, polygenic score.

#### Assortative mating

Table 4 demonstrates phenotypic and genotypic correlations for all parental spouse pairs with data in the ALSPAC cohort. Phenotypic spousal correlations are positive for all phenotypes, and similar to those estimated in other studies (cf. 0.41^39^, 0.62^55^ and 0.66^56^). This provides evidence of phenotypic assortative mating on both educational attainment and socioeconomic position between ALSPAC parents. To test if this phenotypic sorting induces genetic correlations between spouses, we examined genetic correlations between spouses based upon a polygenic score built from all SNPs and a polygenic score that was restricted to SNPs associated with education at genome-wide levels of significance (p-value<5E^-08^).^14^ Positive genetic correlations were observed between spouse-pairs for both education polygenic scores, suggesting that the observed phenotypic assortment may have induced genetic assortment. The presence of education assortative mating in the parents’ generation of our data will likely to upwards bias in heritability estimates of educational attainment and therefore also inflate genetic correlation estimates between education and other phenotypes. The spousal phenotypic correlation for CRP was 0.004 (0.030), and the spousal correlation of the CRP polygenic score was - 0.009 (0.027). These results contrast to the spousal correlations on the social variables and imply no assortment on CRP.

**Table 4:** Correlations between spouses.

|  | Spouses | n |
| --- | --- | --- |
| *Phenotype* |  |  |
| Highest educational attainment | 0.560 (0.011) | 5,353 |
| Linear socioeconomic position | 0.434 (0.010) | 6,858 |
| Binary socioeconomic position | 0.297 (0.008) | 5,737 |
| CRP | 0.004 (0.030) | 1,129 |
| *Genotype* |  |  |
| Education genetic score all SNPs | 0.181 (0.025) | 1,262 |
| Education genetic score GWAS SNPs | 0.080 (0.027) | 1,262 |
| CRP genetic score GWAS SNPs | -0.009 (0.027) | 1,385 |

To further explore the potential impact of assortative mating on our results we conducted additional sensitivity analyses in GCTA controlling for a parent’s years of education and socioeconomic position. This method still assumes no assortative mating, but inconsistency between the main results and these sensitivity analyses provide an indication of heritability inflation due to assortative mating. The results of these analyses (Figure 4) demonstrate that the heritability of educational attainment at age 16 is greatly attenuated – at around half the estimated effect size - when parental education or socioeconomic position is controlled for. This suggests that differences in educational attainment which are associated with common genetic variation can in part be explained by assortative mating on parental education and socioeconomic position. When these sensitivity analyses were applied to genetic correlation estimates between education and socioeconomic position, the impact of assortative mating was less clear due to greater estimation imprecision (Figure S1 in supplementary material).

**Figure 4:**
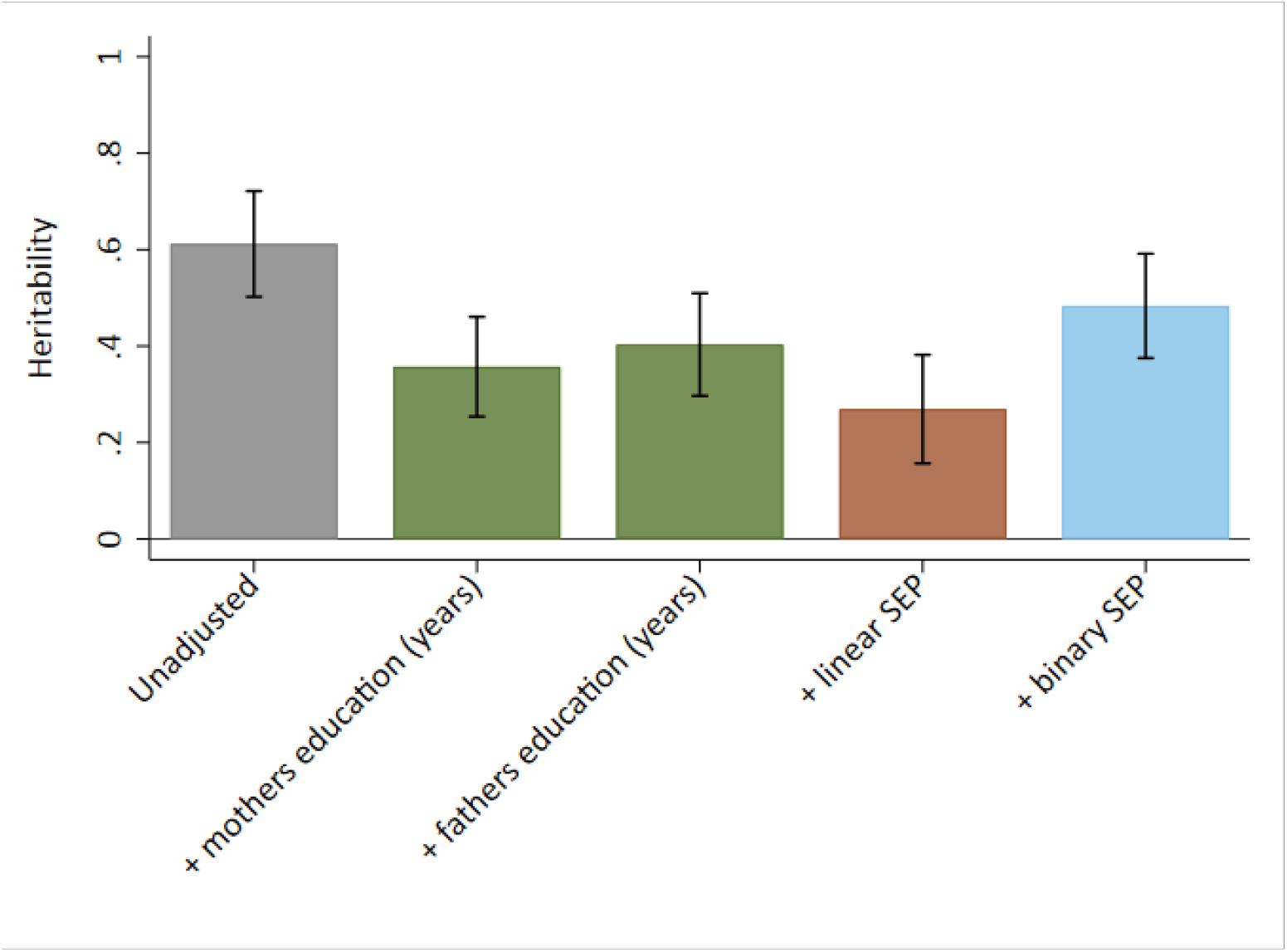
Heritability of educational attainment adjusting for parental socioeconomic variables. Grey bar represents the estimated heritability of educational attainment measured at age 16; green bars represent heritability adjusted for mothers and fathers’ years of education; orange bar represents heritability adjusted for linear socioeconomic position; blue bar represents heritability adjusted for binary socioeconomic position.

#### Negative confounding

Negative confounding, where an environmental factor operates in the opposite direction of effect on two phenotypes, can inflate bivariate heritability estimates. For example, suppose that higher SEP families choose to live in areas with lower air quality (e.g. cities and other urban areas), which results in adverse child neurodevelopment^57^. In this scenario, the negative environmental effect of location on educational attainment operates in the opposite direction to the positive effect of SEP on educational attainment (Figure 5). Negative confounding could also result from positive discrimination in educational settings whereby children are treated differently depending on their social background. Negative confounding leads to artificial reduction of phenotypic correlations which can result in genetic correlation explaining more than 100% of the phenotypic association. To investigate if negative confounding could produce bivariate heritabilities greater than one, we ran a series of simulations (see supplementary material for the data generating process). We defined the variables according to the variation they explained in two phenotypes *A* and *B* (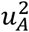 and 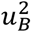 respectively) and the correlation between these effects (r_u_). Figure 6 illustrates the range of values of 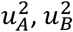 and *r_u_* that were required to produce a bivariate heritability between educational attainment at age 16 and measures of SEP greater than one. This pattern and strength of negative confounding is consistent across all ages of education and both measures of SEP (Figures S2-S3 in supplementary material). The simulations demonstrate that negative confounding is a possible driver of the high bivariate heritabilities we observe where the confounder explains a high amount of variance in either phenotype, if there is a strong correlation between its effects on the phenotypes, or a more moderate function of both.

**Figure 5:**
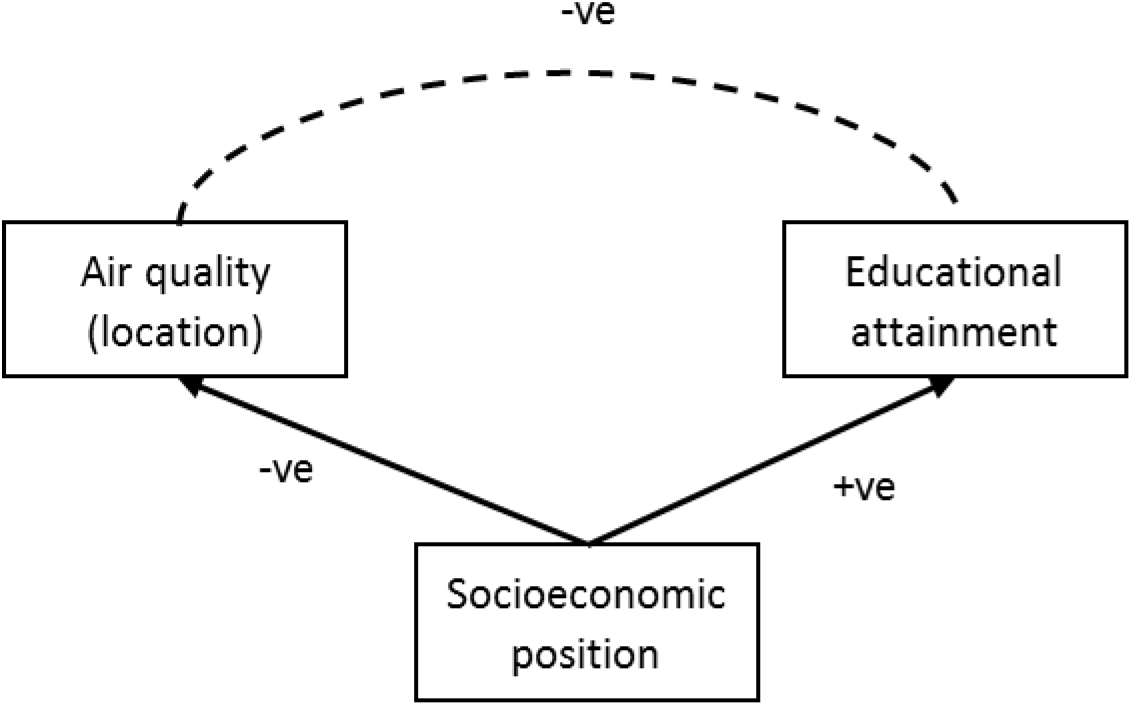
Negative confounding caused by socioeconomic position. In this scenario socioeconomic position has a negative association with air quality and a positive association with educational attainment and, which induces a spurious negative correlation between air quality and educational attainment.

**Figure 6:**
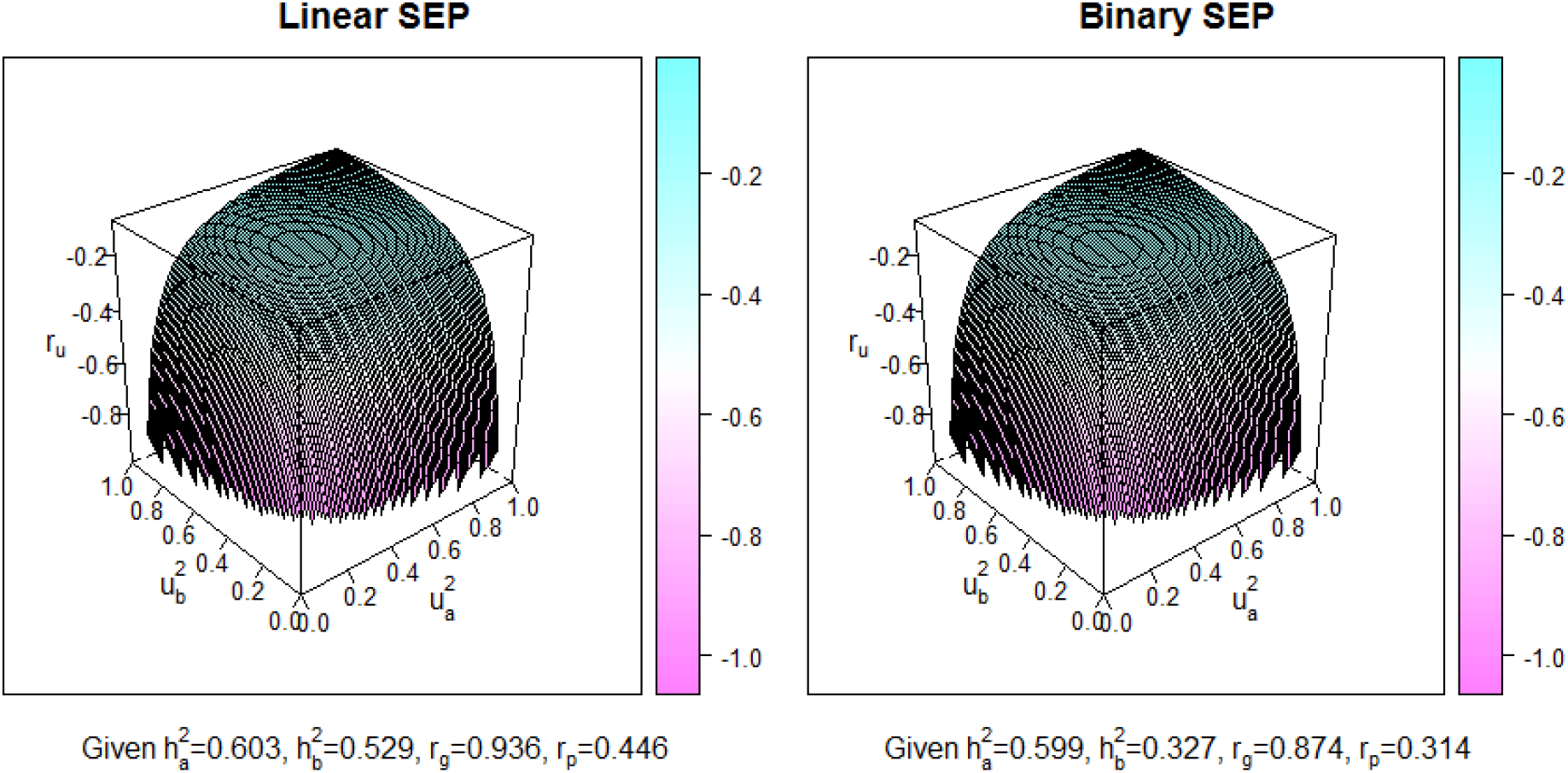
Negative confounding required to obtain a bivariate heritability greater than one for educational attainment at age 16 and socioeconomic position. 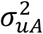, heritability of negative confounder for phenotype ***A***; 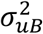, heritability of negative confounder for phenotype ***B; r_u_***, correlation between effects of confounder on phenotypes ***A*** and ***B***.

## Discussion

These results demonstrated how structural and environmental mechanisms can induce bias in estimates of genetic contributions to phenotypes from samples of unrelated individuals. The presence of genetic association does not necessarily imply biologically entrenched contributions and may reflect environmental mechanisms such as population stratification, assortative mating and dynastic effects. Violations to the assumptions made by methods such as GCTA-GREML may lead to inflated estimates of heritability and genetic correlation. Our results add to the growing body of evidence that these methods may overestimate heritability and build upon recently developed methods for estimating genetic associations in the presence of confounding biases.^34,37,38^ Awareness of these issues is also important for interpreting Mendelian randomization studies.^47^ Social phenotypes such as educational attainment and SEP, which are complex, highly assortative and dynastic, appear to be particularly susceptible to these factors. It is therefore important that studies within the rapidly growing area of sociogenomic research^58^ are account for these biases. These results are important as they demonstrate that analyses using samples of unrelated individuals may not provide estimates of heritability or genetic correlation that are driven solely by causal biological factors. Educational attainment and SEP may be influenced by shared genetic effects, and it is therefore important that these potential sources of bias are well understood.

Our SNP heritability estimates of educational attainment were broadly similar to those estimated from twin studies,^18–21^ and there was evidence of increasing heritability with age as has been previously observed.^59^ SNP heritability is expected to be estimated as lower than twin heritability because it only accounts for the additive genetic variance of common SNPs, whereas twin heritability accounts for all genetic variation including rare variants, dominance and epistasis. A recently developed method using whole genome sequence data has demonstrated that a large proportion of SNP heritability may be captured by rare variants, though this was only demonstrated for height and BMI.^60^ Previous studies have highlighted that GREML heritability estimates are upwards biased by family effects^34,37^, and this bias may be present in our results. However, the strength of this bias may be smaller for education test scores which are likely to capture a more cognitive aspect of educational performance than the more social aspect of education that years of education captures. As we have discussed in more detail elsewhere,^61^ the high heritabilities we observed may also reflect genuine differences due to the spatiotemporal homogeneity of the ALSPAC cohort. The mechanisms we investigated may also have larger effects in the ALSPAC study as a regional cohort than in other data samples; the impact of these mechanisms on more geographically dispersed studies such as UK Biobank is currently unknown. Our estimates of the SNP heritability of cognitive ability (41.7%) and SEP (linear: 54.7%; binary: 33.7%) were broadly similar to educational attainment, and also exceeded those in previous studies of 29% for cognitive ability^12^ and 20% for SEP.^11–13^ That heritability was higher in the linear measure than the binary measure of SEP may reflect our cut-point in determining ‘high’ vs ‘low’ for the binary classification or genuine differences between the two measures. The estimates of proportion of variation in all phenotypes explained by the educational attainment polygenic score was broadly consistent with previous research^3^ which used an earlier version of the EA score.^15^ Estimated genetic correlations between educational attainment, SEP and cognitive ability were consistent with findings from other cohorts^5,11,12^ but with greater statistical precision due to larger sample sizes and the precision of GCTA over other methods.^25^ That genetic correlations between educational attainment and cognitive ability persist so strongly throughout childhood further demonstrates the importance of cognitive ability to education in the UK.^8^ Further research is required to investigate how these genetic relationships persist into further and higher education. Bivariate heritability analyses indicated that shared genetic variation can explain a substantial fraction of the phenotypic correlations between educational attainment, SEP and cognitive ability. However, our simulations demonstrate that the contribution of genetic variation to education may be overestimated due to negative confounding.

Attenuation of genetic associations between children’s polygenic score and educational attainment was between one and two thirds after controlling for both parents polygenic scores, supporting the presence of dynastic effects whereby parental genotype indirectly effected offspring phenotype. Furthermore, both parents’ scores remained robust predictors of children’s attainment over and above the child’s polygenic score. Phenotypic spousal correlations demonstrated strong evidence of parental assortative mating on educational attainment (r=0.56) and SEP (r=0.43), which induced genetic correlations at education associated loci of r =0.18. Heritability estimates of educational attainment were attenuated by roughly half when parental education or SEP was controlled for. This supports heritability inflation due to assortative mating and/or dynastic effects in ALSPAC. We found no strong evidence that our estimates were biased by population stratification as measured by the genetic principal components, but this may reflect the inability of genetic principle components to capture subtle population structure^62^ rather than adequate control of it.^23,43^ It is also possible that our high estimates reflected the relatively homogenous educational environment experienced by the ALSPAC cohort when compared to previous studies. Environmental homogeneity increases the proportion of variation that can be attributed to genetic effects, and the ALSPAC children were all born within three years and mostly experienced the same school system within the same region of the UK. Our negative control analyses provided little evidence of dynastic effects or assortative mating for CRP in our sample. While this is expected, it strengthens confidence that the dynastic effects and assortative mating we observe for educational attainment are robust and do not arise from other biasing factors such as genotyping errors.

Several limitations must be acknowledged in this study. First, measurement error on the phenotypes may have influenced our results. Genotyping accuracy and strict quality controls on the genetic data and educational attainment taken from administrative records should result in insufficient measurement error in these phenotypes to meaningfully bias our estimates. However, there may be some measurement inaccuracy in how well the education test scores capture underlying educational ability over and above test-retest reliability. Measurement error will be greater on SEP as these measures relied on self-reported data, but this would have to be differential and patterned to bias estimates (independent non-differential measurement error will only reduce statistical precision of the estimates, not bias them). Second, uneven linkage disequilibrium and further residual population structure in the ALSPAC genetic relatedness matrix could bias our results.^34,37,63,64^ We controlled for the first twenty principle components of population structure in our full analyses, but this is unlikely to account for all differences. Another possible source of confounding in our study is that of shared environmental factors^65^ due to schooling. Within our sample many children will attend the same schools and because educational attainment and SEP are both associated with schooling in the UK, this could inflate heritability through its attribution to additive genetic variation. Recent research has demonstrated the importance of geography as a source of bias in heritability studies^66^ and because we use a heavily geographically clustered cohort this may inflate our heritability estimates. Third, the definition of educational attainment used in the GWAS to conduct the polygenic score was years of education, which is relatively crude and does not discriminate academic performance within each additional year of education. It is therefore possible that the score we use is capturing a social rather than performance aspect of education. Finally, it is possible that our estimates could have been inflated by cryptic relatedness. To overcome this, we restricted our analytical sample to individuals with identity by descent less than 0.1, but it remains possible that some related participants (closer than 1^st^ cousin) will have been included. While data on mother-father-offspring trios provide opportunities to investigate the presence and strength of these biases, mother-father-offspring-sibling quad approaches may offer further opportunities to test for heterogeneity in dynastic effects between siblings.

In conclusion, our results demonstrate some of the causal structures that may bias univariate and bivariate genetic estimates such as heritability and genetic correlations, particularly when applied to complex social phenotypes. Future studies may make use of the methodological tools that we highlight here to assess these alongside others.^34,37,38^ Principally, studies based upon within-family or between-sibling designs will be better equipped to provide informative and unbiased genetic associations given their robustness to population stratification, dynastic effects and assortative mating. Genetic studies investigating complex social relationships should be interpreted with care in light of these mechanisms.

## Data and methods

### Study sample

Participants were children from the Avon Longitudinal Study of Parents and Children (ALSPAC). Pregnant women resident in Avon, UK with expected dates of delivery 1st April 1991 to 31^st^ December 1992 were invited to take part in the study. The initial number of pregnancies enrolled was 14,541. When the oldest children were approximately 7 years of age, an attempt was made to bolster the initial sample with eligible cases who had failed to join the study originally. This additional recruitment resulted in a total sample of 15,454 pregnancies, resulting in 14,901 children who were alive at one year of age. From these there is genetic data available for 7,748 children on at least one of educational attainment, socioeconomic position and cognitive ability after quality control and removal of related individuals (see *Genetic data* below). We use the largest available samples in each of our analyses to increase precision of estimates, regardless of whether a child contributed data to the other analyses. For full details of the cohort profile and study design see ^67,68^. Please note that the study website contains details of all the data that is available through a fully searchable data dictionary and variable search tool at http://www.bristol.ac.uk/alspac/researchers/our-data/. The ALSPAC cohort is largely representative of the UK population when compared with 1991 Census data; there is under representation of some ethnic minorities, single parent families, and those living in rented accommodation.^67^ Ethical approval for the study was obtained from the ALSPAC Ethics and Law Committee and the Local Research Ethics Committees. Consent for biological samples has been collected in accordance with the Human Tissue Act (2004).

### Genetic data

DNA of the ALSPAC children was extracted from blood, cell line and mouthwash samples, then genotyped using references panels and subjected to standard quality control approaches. ALSPAC children were genotyped using the Illumina HumanHap550 quad chip genotyping platforms by 23andme subcontracting the Wellcome Trust Sanger Institute, Cambridge, UK and the Laboratory Corporation of America, Burlington, NC, US. ALSPAC mothers were genotyped using the Illumina human660W-quad array at Centre National de Génotypage (CNG) and genotypes were called with Illumina GenomeStudio. ALSPAC fathers and some additional mothers were genotyped using the Illumina HumanCoreExome chip genotyping platforms by the ALSPAC lab and called using GenomeStudio. All resulting raw genome-wide data were subjected to standard quality control methods in PLINK (v1.07). Individuals were excluded on the basis of gender mismatches; minimal or excessive heterozygosity; disproportionate levels of individual missingness (>3%) and insufficient sample replication (IBD < 0.8). Population stratification was assessed by multidimensional scaling analysis and compared with Hapmap II (release 22) European descent (CEU), Han Chinese, Japanese and Yoruba reference populations; all individuals with non-European ancestry were removed. SNPs with a minor allele frequency of < 1%, a call rate of < 95% or evidence for violations of Hardy-Weinberg equilibrium (P < 5E-7) were removed. Cryptic relatedness was assessed using a IBD estimate of more than 0.125 which is expected to correspond to roughly 12.5% alleles shared IBD or a relatedness at the first cousin level. Related participants that passed all other quality control thresholds were retained during subsequent phasing and imputation. For the mother’s, samples were removed where they had indeterminate X chromosome heterozygosity or extreme autosomal heterozygosity. After quality control, 9,115 participants and 500,527 SNPs for the children, 9,048 participants and 526,688 SNPs for the mothers, and 2,201 participants and 507,586 SNPs for the fathers (and additional mothers) passed these quality control filters.

We combined 477,482 SNP genotypes in common between the samples. We removed SNPs with genotype missingness above 1% due to poor quality and removed participants with potential ID mismatches. This resulted in a dataset of 20,043 participants containing 465,740 SNPs (112 were removed during liftover and 234 were out of HWE after combination). We estimated haplotypes using ShapeIT (v2.r644) which utilises relatedness during phasing. The phased haplotypes were then imputed to the Haplotype Reference Consortium (HRCr1.1, 2016) panel of approximately 31,000 phased whole genomes. The HRC panel was phased using ShapeIt v2, and the imputation was performed using the Michigan imputation server. This gave 8,237 eligible children, 8,675 eligible mothers and 1,722 eligible fathers with available genotype data after exclusion of related participants using cryptic relatedness measures described previously.

### Educational attainment

We use average fine graded point scores at the three major Key Stages of education in the UK at ages 11, 14 and 16. Point scores were obtained from the Key Stage 4 (age 16) database of the UK National Pupil Database (NPD) through data linkage to the ALSPAC cohort. The NPD represents the most accurate record of individual educational attainment available in the UK. The Key Stage 4 database provides a larger sample size than the earlier two Key Stage databases and contains data for each. Fine graded point scores provide a richer measure of a child’s attainment than level bandings and were therefore chosen as the most accurate method of determining academic attainment during compulsory schooling.

### Parental socioeconomic position

We use two measures of parental socioeconomic position; a binary classification based on the widely used Social Class based on Occupation (formerly Registrar Generals Social Class) of “high” [I and II] vs “low” [III-Non-manual, III-Manual, IV, and V] social classes and a continuous classification based upon the Cambridge Social Stratification Score (CAMSIS). Social Class based on Occupation assumes within-strata social homogeneity with clear boundaries, while CAMSIS provides a more flexible measure that accounts for social heterogeneity.^69^

### Cognitive ability

Cognitive ability was measured during the direct assessment at age eight using the short form Wechsler Intelligence Scale for Children (WISC) from verbal, performance, and digit span tests^70^ and administered by members of the ALSPAC psychology team overseen by an expert in psychometric testing. The short form tests have high reliability^71^ and the ALSPAC measures utilise subtests with reliability ranging from 0.70 to 0.96. Raw scores were recalculated to be comparable to those that would have been obtained had the full test been administered and then age-scaled to give a total overall score combined from the performance and verbal subscales.

### Educational attainment polygenic score

To test dynastic effects we used an educational attainment polygenic score with the 1,271 independent SNPs identified to associate with years of education at genome-wide levels of significance (p<5e^-8^) in GWAS^14^ using the software package PRSice.^72^ PRSice was used to thin SNPs according to linkage disequilibrium through clumping, where the SNP with the smallest *P*-value in each 250kb window was retained and all other SNPs in linkage disequilibrium with an r^2^ of >0.1 were removed). This score was generated using GWAS results which had removed ALSPAC and 23andMe participants from the meta-analysis. In the GWAS the score using the 1,271 genome-wide significant SNPs explained 2.5-3.8% of the variation in educational attainment in the two prediction cohorts.

### Negative control analyses

To ensure that our analyses were correctly identifying the biasing mechanisms that we outline and did not represent other biasing factors such as genotyping errors, we ran sensitivity analyses using C-reactive protein (CRP) as a negative control phenotype. CRP is a biomarker of inflammation that is associated with a range of complex diseases and is unlikely to be influenced by the biases that relate to the social phenotypes that we investigate. Systematic population differences (population stratification) in CRP have been found to be insubstantial;^73,74^ parental phenotypic expression of CRP is unlikely to influence offspring CRP (dynastic effects); and parents are very unlikely to selectively mate based upon CRP (assortative mating). Offspring CRP was measured from non-fasting blood assays taken during direct assessment when the children were aged 9.

### Statistical analysis

We estimate SNP heritability (hereafter referred to as heritability) using generalized restricted maximum likelihood (GREML) in the software package GCTA.^22^ GCTA uses measured SNP level variation across the whole genome to estimate the proportion of variation in educational attainment, socioeconomic position, and cognitive ability that can be explained by common genetic variation. We use a series of univariate analyses of the form:

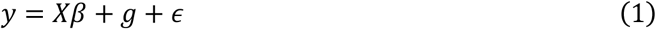

where *y* is the phenotype, *X* is a series of covariates, *g* is a normally distributed random effect with variance 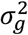, and *ϵ* is residual error with variance 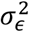. Heritability is defined as the proportion of total phenotypic variance (genetic variance plus residual variance) explained by common genetic variation:

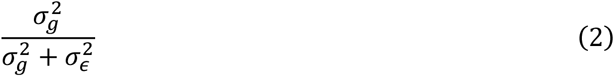

Where genetically similar pairs are more phenotypically similar than genetically dissimilar pairs heritability estimates will be non-zero.

We estimate genetic correlations using bivariate GCTA, running nine sets of analyses between educational attainment at each age and each of linear socioeconomic position, binary socioeconomic position, and cognitive ability. Genetic correlation quantifies the extent to which genetic variants that associate with one phenotype (i.e. educational attainment) also associate with another phenotype (i.e. cognitive ability). It therefore refers to the correlation of all genetic effects across the genome for phenotypes *A* and *B,* and is estimated as:

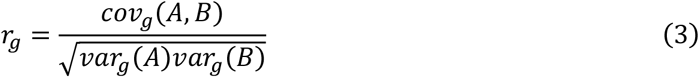

Where *r_g_* is the genetic correlation between phenotypes *A* and *B, var_g_*(*A*) is the genetic variance of phenotype *A* and *cov_g_*(*A,B*) is the genetic covariance between phenotypes *A* and *B*. Genetic correlations can indicate that two phenotypes are influenced by the same genetic variants (i.e. have shared genetic architecture). In contrast, the bivariate heritability is the proportion of the phenotypic correlation that can be explained by the genotypes. Genetic correlations and bivariate heritability are likely to differ. For example, two phenotypes may be highly genetically correlated, but if they have low heritability then the bivariate heritability will be low. Bivariate heritability estimates the proportion of phenotypic correlations that can be explained by genetics. It is estimated as:

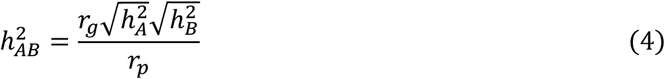

Where *r_g_* is the genetic correlation between phenotypes *A* and *B*, 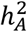 and 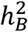 are the heritabilities of phenotype *A* and *B* respectively, and *r_p_* is the phenotypic correlation between phenotypes *A* and B. We used data for unrelated participants (less related than 2^nd^ cousins), as indicated by the ALSPAC Genetic Relatedness Matrices (GRM). Population stratification is controlled for by using the first 20 principal components of inferred population structure as covariates in analyses. Continuous variables were inverse normally transformed to have a normal distribution, a requirement of GCTA.

## Acknowledgements

We are extremely grateful to all the families who took part in this study, the midwives for their help in recruiting them, and the whole ALSPAC team, which includes interviewers, computer and laboratory technicians, clerical workers, research scientists, volunteers, managers, receptionists and nurses. The UK Medical Research Council and Wellcome (Grant ref: 102215/2/13/2) and the University of Bristol provide core support for ALSPAC. A comprehensive list of grants funding is available on the ALSPAC website (http://www.bristol.ac.uk/alspac/external/documents/grant-acknowledgements.pdf); data used at age 23 was specifically funded by the Wellcome Trust and MRC [102215/2/13/2]. Study data were collected and managed using REDCap electronic data capture tools hosted at the University of Bristol. REDCap (Research Electronic Data Capture) is a secure, web-based application designed to support data capture for research studies, providing 1) an intuitive interface for validated data entry; 2) audit trails for tracking data manipulation and export procedures; 3) automated export procedures for seamless data downloads to common statistical packages; and 4) procedures for importing data from external sources. GWAS data was generated by Sample Logistics and Genotyping Facilities at Wellcome Sanger Institute and LabCorp (Laboratory Corporation of America) using support from 23andMe. The Economics and Social Research Council (ESRC) support NMD via a Future Research Leaders grant [ES/N000757/1] and TTM via a postdoctoral research fellowship [ES/S011021/1]. The Medical Research Council (MRC) and the University of Bristol support the MRC Integrative Epidemiology Unit [MC_UU_12013/1, MC_UU_12013/9, MC_UU_00011/1]. This publication is the work of the authors and Tim Morris will serve as guarantor for the contents of this paper. No funding body has influenced data collection, analysis or its interpretations. This work was carried out using the computational facilities of the Advanced Computing Research Centre - http://www.bris.ac.uk/acrc/ and the Research Data Storage Facility of the University of Bristol - http://www.bris.ac.uk/acrc/storage/. This research was conducted using the UK Biobank Resource.

## Footnotes

i The reported genetic correlation of 1.02 is likely due to estimation imprecision.

ii Note that the upper confidence interval represents the upper bound of genetic correlation based upon the estimates, but in practice genetic correlation of would be constrained to the value of 1.

iii Using the values of *r_g_* = 0.965; 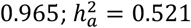; 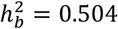; *r_p_* = 0.720, calculated as: 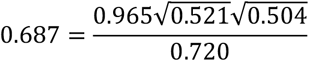

