## Supplementary material for "Why are education, socioeconomic position and intelligence genetically correlated?"

**Figure S1: Genetic correlations between education and socioeconomic position.** Left panel: genetic correlations between educational attainment and linear socioeconomic position. Right panel: genetic correlations between educational attainment and binary socioeconomic position.

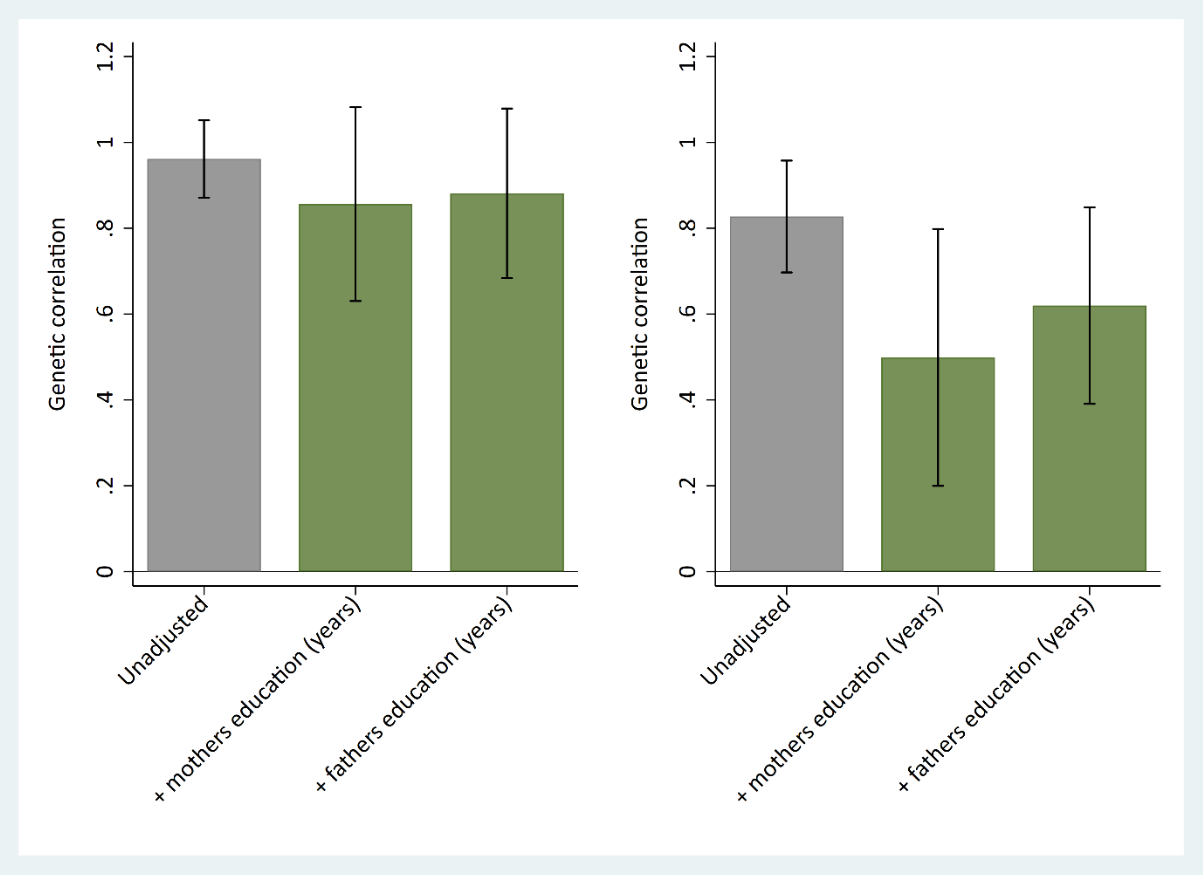

### Negative confounding

Bivariate heritabilities describe the proportion of phenotypic similarity between two traits that is due to genetic similarity. As such, the values of bivariate heritability are expected to be constrained to values within the range of zero to one, where zero indicates no genetic influence on phenotypic similarity and one indicates total genetic influence on phenotypic similarity. One possibility for estimates of bivariate heritability greater than one is the presence of negative confounding, whereby a secondary variable acts in opposite directions on the two traits. To explore this possibility, we ran a series of simulations to determine the amount of negative confounding that would be required to obtain bivariate heritabilities greater than one given our observed trait heritabilities, genetic correlations and phenotypic correlations.

The model underlying these simulations is as follows:

$$y_A = g_A + u_A + e_A$$

$$y_B = g_B + u_B + e_B$$

$$g_A \sim g_B \sim MVN(0, \Sigma_g^2)$$

$$u_A \sim u_B \sim MVN(0, \Sigma_u^2)$$

$$e_A \sim N(0, 1 - \sigma_{gA}^2 - \sigma_{uA}^2)$$

$$e_B \sim N(0, 1 - \sigma_{gB}^2 - \sigma_{uB}^2)$$

$$\Sigma_g^2 = \begin{bmatrix} \sigma_{gA}^2 & \rho_g \\ \rho_g & \sigma_{gB}^2 \end{bmatrix}$$

$$\Sigma_u^2 = \begin{bmatrix} \sigma_{uA}^2 & \rho_u \\ \rho_u & \sigma_{uB}^2 \end{bmatrix}$$

Where $y_A$ is phenotype A, $y_B$ is phenotype B, $g_A$ and $g_B$ denote genetic effects for phenotypes A and B drawn from a multivariate normal distribution with variance $\Sigma_g^2$, $u_A$ and $u_B$ denote the effects of a confounder variable $u$ on phenotypes A and B drawn from a multivariate normal distribution with variance $\Sigma_u^2$, and $e_A$ and $e_B$ denote normally distributed residuals for phenotypes A and B. The confounder is specified in two ways:

1. With respect to the proportion of variance that it explains in phenotypes $y_A$ and $y_B$ ($\sigma_{uA}^2$ and $\sigma_{uB}^2$ respectively)
2. The correlation between $u_a$ and $u_b$ ($\rho_u$)

The full range of values (0:1) that $\sigma_{uA}^2$, $\sigma_{uB}^2$ and $\rho_u$ can take to satisfy a bivariate heritability estimate that is greater than one, given the observed heritabilities, genetic correlation and phenotypic correlations, can therefore be computed and plotted for each pair of traits. Figures 1-3 below plot the full range of values computed from R code for generating these scenarios (available here).

Figure S2: Negative confounding required to obtain a bivariate heritability greater than one for educational attainment at age 11 and socioeconomic position. $\sigma^2_{uA}$, heritability of negative confounder for phenotype $A$; $\sigma^2_{uB}$, heritability of negative confounder for phenotype $B$; $r_u$, correlation between effects of confounder on phenotypes $A$ and $B$.

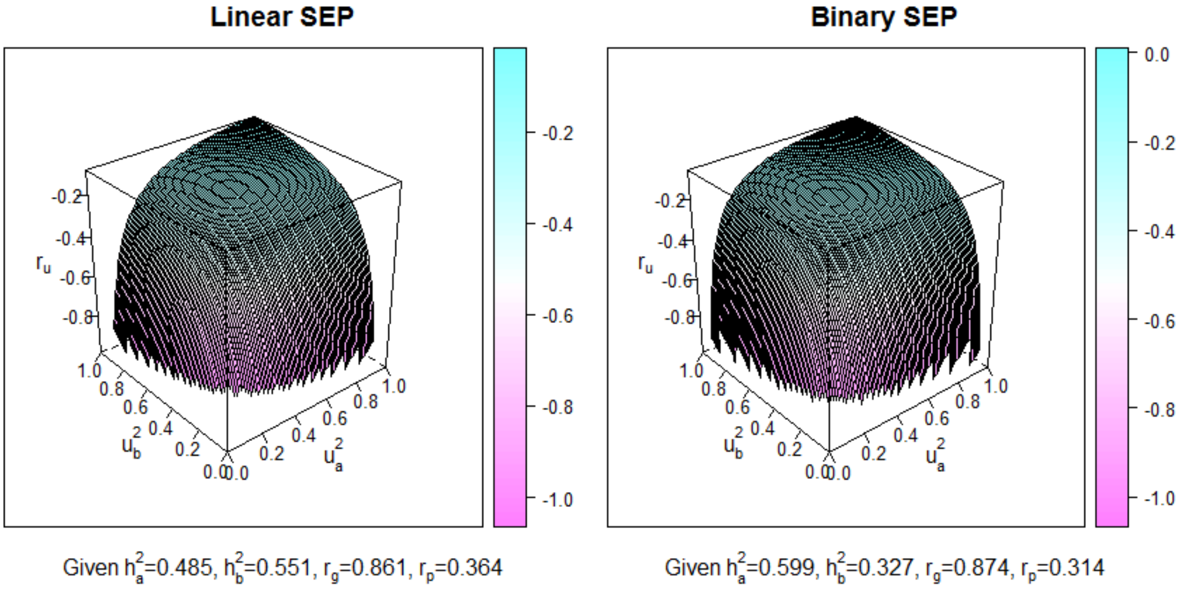


Figure S3: Negative confounding required to obtain a bivariate heritability greater than one for educational attainment at age 14 and socioeconomic position. $\sigma^2_{uA}$, heritability of negative confounder for phenotype $A$; $\sigma^2_{uB}$, heritability of negative confounder for phenotype $B$; $r_u$, correlation between effects of confounder on phenotypes $A$ and $B$.

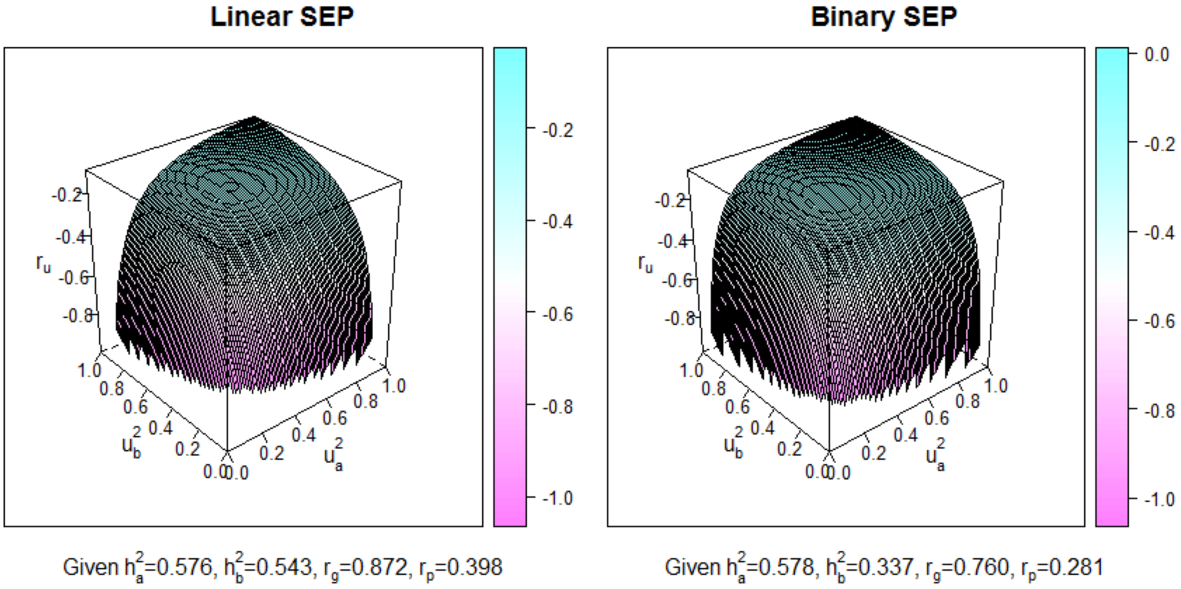
